# Benchmarking of the Oxford Nanopore MinION sequencing for quantitative and qualitative assessment of cDNA populations

**DOI:** 10.1101/048074

**Authors:** Spyros Oikonomopoulos, Yu Chang Wang, Haig Djambazian, Dunarel Badescu, Jiannis Ragoussis

## Abstract

To assess the performance of the Oxford Nanopore Technologies MinION sequencing platform, cDNAs from the External RNA Controls Consortium (ERCC) RNA Spike-In mix were sequenced. This mix mimics mammalian mRNA species and consists of 92 polyadenylated transcripts with known concentration. cDNA libraries were generated using a template switching protocol to facilitate the direct comparison between different sequencing platforms. The MinION performance was assessed for its ability to sequence the cDNAs directly with good accuracy in terms of abundance and full length. The abundance of the ERCC cDNA molecules sequenced by MinION agreed with their expected concentration. No length or GC content bias was observed. The majority of cDNAs were sequenced as full length. Additionally, a complex cDNA population derived from a human HEK-293 cell line was sequenced on an Illumina HiSeq 2500, PacBio RS II and ONT MinION platforms. We observed that there was a good agreement in the measured cDNA abundance between PacBio RS II and ONT MinION (r_spearman_=0.57, fragments with length more than 700bp) and between Illumina HiSeq 2500 and ONT MinION (r_spearman_=0.61). This indicates that the ONT MinION can sequence quantitatively both long and short full length cDNA molecules.

## Introduction

Transcriptome sequencing using short read technologies (Illumina HiSeq 2500^1^, Ion Proton^2^) provides valuable information on transcript abundance, rare transcripts and variable transcription start or end sites. Nevertheless, inferring alternatively spliced isoforms of genes from short read data through statistical assignment of the most probable combination of exons is still computationally challenging and not very accurate^3^. Uneven read coverage^4^, complex splicing^5^ and potential sequencing bias^6^ complicates even more the task. The PacBio RS II^7^ zero–mode waveguide (ZMW) long-read sequencing technology has proven capable of characterizing the transcriptome in its native, full-length form unravelling novel gene isoforms not previously observed in RNA-seq experiments^8^. Recently, another long-read DNA sequencing technology based on nanopore sequencing (MinION) was introduced from Oxford Nanopore Technologies Ltd (ONT)^9^. Similar to PacBio RS II, the ONT MinION has been shown that can resolve the exon structure of mRNA molecules transcribed from genes with a large variety of isoforms^10^.

Here, we assessed the ONT MinION performance for its ability to sequence the cDNAs as full length and with good accuracy in terms of cDNA abundance and sequence identity. To evaluate the performance of the ONT MinION platform the cDNA of a commercially available defined set of 92 polyadenylated transcripts, that mimic mammalian mRNA species (ERCC RNA Spike-In mix), were sequenced in both an Illumina HiSeq 2500 or MiSeq instruments and ONT MinION. Additionally, a complex cDNA population from a HEK-293 cell line was sequenced in three sequencing platforms (Illumina HiSeq 2500, PacBio RS II, ONT MinION) and the agreement in the cDNA abundance of the different transcript isoforms across the three platforms was assessed. The results indicate that the ONT MinION platform can sequence quantitatively cDNA molecules similar with the Illumina and PacBio RS II platforms paving the way for full length sequencing of cDNA molecules with nanopores.

## Results

### Sequencing the ERCC cDNA molecules on the ONT MinION platform

We sequenced between 9,525 and 197,014 ERCC cDNA molecules in four different ONT MinION flow cells as presented in Table I. We used two different versions of the ONT MinION flowcells an old version r7 and a newer version r7.3. The different types of reads produced from the ONT MinION platform has been described in detail in other studies^11^. Briefly, the ONT MinION platform can sequence both strands of the same DNA molecule at once due to the presence of a hairpin adaptor. The strand that is sequenced first is called the “template read”. The strand that is sequenced second is called the “complement read”. If both strands of the same molecule are sequenced, a consensus sequence (“2D read”) is produced from the “template read” and the “complement read”. After the sequencing, the ONT analysis pipeline separates the sequenced reads in two groups the “pass” and the “failed” group. The “pass” group contains high quality 2D reads (average base quality score of the “2D read” >=9) along with their “template” and “complement” read sequence. The “failed” group contains the rest of the reads. The reads in the “pass” and “fail” group are referred in this manuscript as high and low quality reads respectively. In Table I we see that all of the cDNA molecules were sequenced as template strand. A fraction of the cDNA molecules (7-49% of the total reads) was sequenced as both template and complement strand and eventually some of them (4-39% of the total reads) were assigned as 2D reads. The proportion of sequenced reads assigned to each template, complement and 2D read group is similar to other ONT MinION studies that sequenced amplicons of similar size^10,12^.

**Table 1.** Number of reads sequenced in each MinION experiment. The sequenced reads are separated in the template, complement and 2D read types for both the high and low basecalling quality categories. The four flow cells on which the ERCC cDNA library was sequenced are presented. The flow cell of the HEK-293 cDNA library is also presented.?

| flow cell number | 1 | 2 | 3 | 4 | HEK-293 |
| --- | --- | --- | --- | --- | --- |
| flowcell version | r7 | r7.3 | r7.3 | r7.3 | r7.3 |
| library protocol version | SQK-MAP003 | SQK-MAP004 | SQK-MAP005 | SQK-MAP005 | SQK-MAP005 |
| basecalling version | r7 2D | r7.X 1.12 2D | r7.X 1.16 2D | r7.X 1.16 2D | r7.X 1.16 2D |
| total number of reads | 25502 | 51163 | 9525 | 197014 | 16540 |
| total number of aligned reads | 6623 | 23218 | 3799 | 122646 | 4521 |
| template reads | 25502 | N/A | N/A | N/A | N/A |
| complement reads | 5998 | N/A | N/A | N/A | N/A |
| 2D reads | 1900 | N/A | N/A | N/A | N/A |
| aligned template reads | 6007 | N/A | N/A | N/A | N/A |
| aligned complement reads | 1081 | N/A | N/A | N/A | N/A |
| aligned 2D reads | 1255 | N/A | N/A | N/A | N/A |
| total aligned individual reads | 6623 | N/A | N/A | N/A | N/A |
| number of low quality reads | N/A | 46986 | 9014 | 167859 | 16374 |
| template reads | N/A | 46986 | 9014 | 167859 | 16374 |
| complement reads | N/A | 12905 | 4167 | 60962 | 998 |
| 2D reads | N/A | 10632 | 3227 | 39609 | 541 |
| aligned template reads | N/A | 14819 | 2399 | 84876 | 4198 |
| aligned complement reads | N/A | 4571 | 819 | 20464 | 106 |
| aligned 2D reads | N/A | 7137 | 1386 | 33210 | 228 |
| total aligned low quality reads | N/A | 19103 | 3288 | 93494 | 4374 |
| number of high quality reads | N/A | 4177 | 511 | 29155 | 166 |
| template reads | N/A | 4177 | 511 | 29155 | 166 |
| complement reads | N/A | 4177 | 511 | 29155 | 166 |
| 2D reads | N/A | 4177 | 511 | 29155 | 166 |
| aligned template reads | N/A | 3552 | 498 | 28602 | 108 |
| aligned complement reads | N/A | 3852 | 480 | 27232 | 101 |
| aligned 2D reads | N/A | 4115 | 511 | 29151 | 146 |
| total aligned high quality reads | N/A | 4115 | 511 | 29152 | 147 |

The MinION community is currently mainly focused on the 2D read type from the high quality category. As has already been observed in other studies^11^ the 2D reads accumulation gradually drops over the sequencing time and it is high only in the beginning of the sequencing run (Supplementary Fig. S1A, B). For accurate gene expression analysis a large number of sequenced reads is preferable and to overcome the 2D read limitation we examined whether some other read types (template, complement, 2D reads) from the low quality category can be used for quantitative expression analysis. We assessed which read type is closer to the expected number of ERCC cDNA molecules from both high and low quality categories and how many sequenced reads can be aligned to the reference ERCC database in each category and read type. We eventually selected the read type that had the highest number of reads and that was closer to the expected distribution of ERCC cDNA molecules.

### The abundance of the different ERCC cDNA molecules sequenced from either the Illumina HiSeq 2500/MiSeq platforms or the ONT MinION is comparable

To test whether the ONT MinION performs equally well as the Illumina platforms in estimating the ERCC cDNA abundance, the same full length ERCC cDNA population used in the ONT MinION experiments, was sequenced on an Illumina HiSeq 2500 or MiSeq instruments. To estimate the cDNA abundance from the Illumina sequencing, both the FPKM method and molecular counting of the sequenced 5’ end or 3’ end fragments of the ERCC cDNA molecules^13^ were examined (Supplementary text). The 5’ end molecular counting was selected as more accurate (Supplementary text) and the number of cDNA molecules calculated from both the ONT MinION and Illumina HiSeq 2500/MiSeq was compared.

Initially we compared the agreement between the number of sequenced cDNA molecules from the ONT MinION platform with the number of RNA molecules as provided from the manufacturer (Ambion), for each one of the 92 ERCC species. In Fig. 1A we see that the ONT MinION sequenced reads are as good as the Illumina 5’ end molecular counts (Fig. 1B) in estimating the cDNA abundance (r_p_=0.98 for MinION, r_p_=0.97 for Illumina). We then assessed which ONT MinION read type (template, complement, 2D for both low and high quality categories) is in better agreement with the Illumina molecular counts. The Illumina molecular counts represent our best estimation of the true cDNA abundance. We saw that independently from the read type of the ONT MinION reads used, the agreement for the low quality category is better than the one for the high quality category. The correlation with the Illumina molecular counts for the low quality category is r_p raw values_=0.98-0.99, r_p log10 values_=0.98 (Supplementary Fig. S2A-C and Supplementary Fig. S3A-C respectively). The same correlation for the high quality category is r_p raw values_=0.93-0.94, r_p loq10 values_=0.96-0.97 (Supplementary Fig. S2D-F and Supplementary Fig. S3D-F respectively). When we pooled together the reads from the low and high quality categories, the agreement for each one of the read types is again high (r_p raw values_ =0.96-0.99, r_p log10 val_ues=0.97-0.98, Supplementary Fig. S2G-I, Supplementary Fig. S3G-I). We then assessed which one of the read type categories, from the pooled quality category, is in better agreement with the Illumina cDNA molecular counts. In both the flow cell v7.3 (Supplementary Fig. S2G-I) and flow cell v7 (Supplementary Fig. S4A-C) we see that the template reads gave a higher and more consistent correlation with the expected Illumina cDNA molecular counts than the complement or 2D reads. The correlation values are r_p raw values v7.3_=0.99 and r_p raw values v7_ =0.99 for the template reads, r_p raw values v7.3_=0.96 and r_p raw values v7_ =0.93 for the complement reads, r_p raw values v7.3_=0.99 and r_p raw values v7_ =0.72 for the 2D reads.

**Figure 1.**
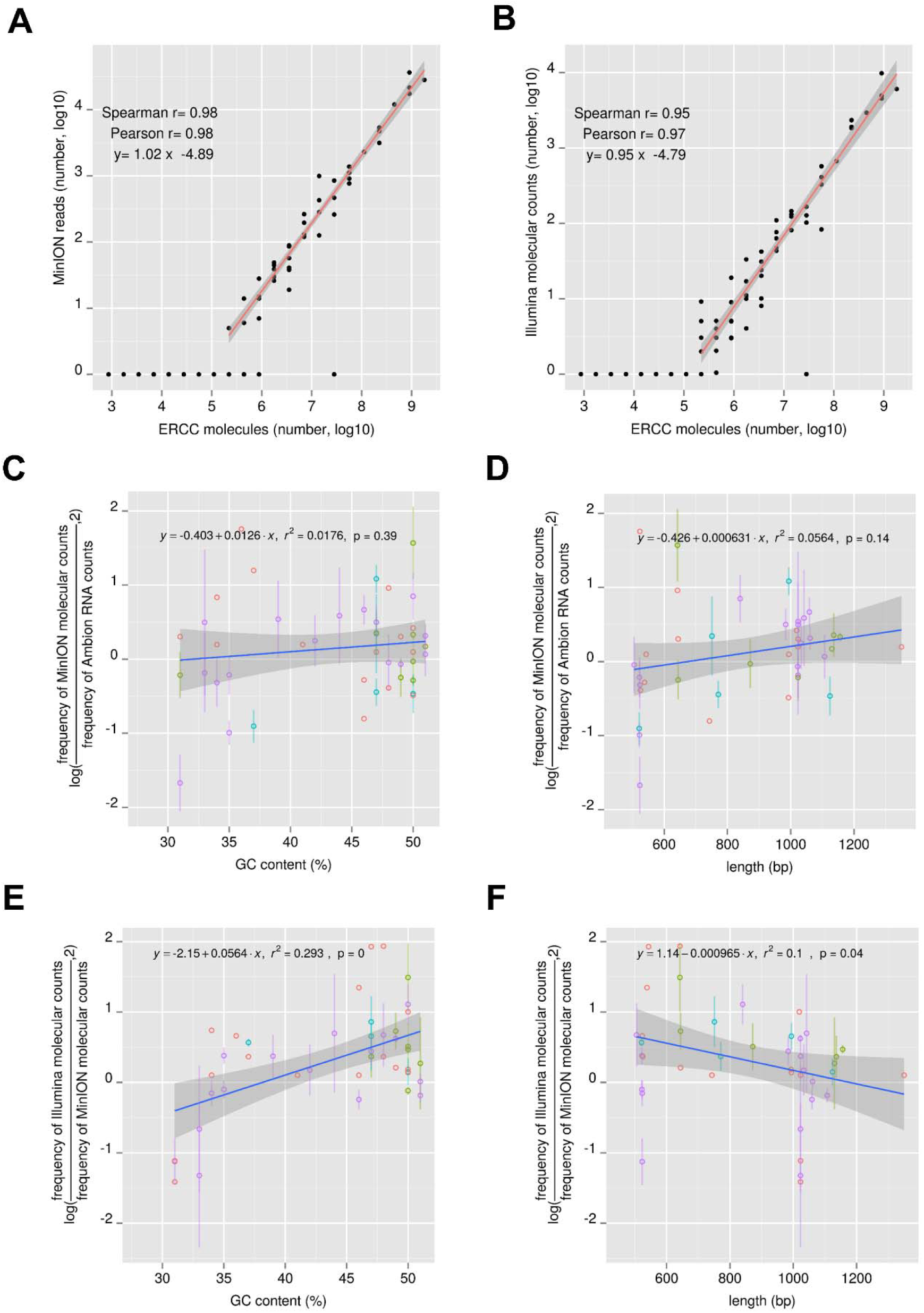
Estimation of the ERCC cDNA abundance with the ONT MinION platform. A, B) Comparison of the ERCC cDNA abundance between the ONT MinION (A) and the Illumina HiSeq 2500 or MiSeq platforms (B) against the expected number of RNA molecules as provided from the manufacturer (Ambion). The template reads from the ERCC MinlON experiments number 1, 2, 3, 4 were pooled together and used for (A). Similarly the corresponding Illumina data were pooled together for (B). The total number of molecules presented on the x-axis corresponds to 3.5 pgs of ERCC RNA. C-F) Deviations of the ERCC expression level estimates with the ONT MinlON platform from the Ambion RNA molecular counts (C,D) or from the Illumina HiSeq 2500/MiSeq estimated cDNA abundance (E,F) as a function of the GC content (C,E) and the ERCC length (D,F). In C-F we plot the log2 ratio of observed (ONT MinION) to expected (Ambion,Illumina) read counts for the ERCC spike-ins (y-axis, log) for each of the samples relative to their length or GC content (x-axis). Due to the variable sequencing depth from each ERCC MinION experiment, each point is the average value from different MinION flow cell runs if at least 5 reads have been detected for this point in the corresponding MinION runs. The points are colored differently based on the number of flow cell runs that they were detected (red, green, cyan, purple correspond to values derived from one, two, three or four flow cell runs respectively). The standard deviation is also presented for points with values from two or more MinION runs.

### The ONT MinION sequenced reads usually correspond to the full length of ERCC cDNA molecules but some sequencing runs can produce partially sequenced molecules

We used the Illumina HiSeq 2500 /MiSeq data to calculate how many of the cDNA molecules are synthesized as full length before sequencing. For this we analyzed the position on the reference ERCC database where the 5’ end fragments align (Supplementary Fig. S5). We saw that 90.8% of them were full length (aligned position of the 5’ Illumina fragments maximum 5 bases from the 5’ position of the reference transcript). We then assessed whether the ONT MinION reads were processed as full length. We separated the ONT MinION reads in two groups. The first group includes reads where the sequencing started from the 5’ end of the transcripts in the ERCC reference database (reads aligning as sense on the database, we call them “sense reads”). The second group includes reads where the sequencing started from the 3’ end (reads aligning as anti-sense on the database, we call them “antisense reads”). We examined what percentage of the sense or antisense reads reached until the 3’ or the 5’ end of the ERCC transcripts respectively. In Supplementary Fig. S6 we see that in all the ERCC runs except run number 4 the reads were sequenced as full length. In the case of run number 4, 77% of the “antisense” template reads (that do not belong in either the low or high quality 2D read type) and 80% of the “sense” template reads are partially processed (the sequencing stops more than 5 bp away from the TES and TSS respectively). Since run number 4 gave the maximum number of reads we used it to assess whether any potential systematic bias during the sequencing run affected the complete processing of reads. For this we examined whether the reads were sequenced as full length over time.

The ERCC library has the advantage that it consists of two abundant groups of reads that can be separated based on their length, a group with short transcripts (average length ~600bp) and a group with long transcripts (average length ~1200 bp). This permits to assess whether the ONT MinION platform preferentially sequences short or long DNA molecules as is the case for the PacBio RS II platform^8^. Additionally it permits to assess whether the performance of the ONT MinION platform is constant throughout the sequencing run. In Supplementary Fig. S7 we see that in the beginning of every restart of the sequencing run (“remux” time^11^) (time point zero, and time points at the purple vertical line) the median sequenced length (~650 bp) was approximately the same (Supplementary Fig. S7B) and the group with the short transcripts was sequenced equally well as the group with the long transcripts. Nevertheless every time there was an increase in the voltage, which drives the DNA translocation across the pore (“bias voltage”^11^), by 5mV (time point at green vertical lines which corresponded to every 4 hours of a continuous sequencing run^11^) we observed a decrease of the median sequenced length by 50-250 bp (Supplementary Fig. S7B). The drop of the median sequenced length followed the drop of the median alignment length on the reference database (from 95% down to 55-85% depending on the time point) (Supplementary Fig. S7D).

Because the sequenced length drop affected both DNA molecules in the short and long transcript group, the overall decrease did not affect the relative quantification of the transcripts. As can be seen in Supplementary Fig. S7E the group with the short transcripts was always more abundant than the group with the long transcripts, independently of the time point during the flow cell run when the reads were produced.

### The ONT MinION platform can sequence equally well short and long ERCC cDNA molecules and shows no specific preference for high or low GC content ERCC cDNAs

We examined whether the ONT MinION platform shows a preference for sequencing better either high or low GC content cDNA molecules and either short or long cDNA molecules. Initially we examined whether any deviation (log fold difference) between the observed ERCC cDNA abundance from the ONT MinION platform and the expected ERCC RNA/cDNA abundance, from either the Ambion molecular counts (RNA concentration) or the Illumina 5’ end molecular counts, is linearly correlated (positively or negatively) with the GC content. The comparison between the ONT MinION cDNA counts and the Ambion RNA counts showed no statistically significant trend (Fig. 1C). On the contrary when we compared the ONT MinION cDNA counts and the Illumina cDNA molecular counts we saw a statistically significant positive trend (r^2^=0.293) (Fig. 1E). This indicates that 29% of the variation in the deviation between the ONT MinION cDNA counts and the Illumina cDNA counts can be attributed to the difference in the GC content of the ERCC cDNA molecules. Consequently, the high GC content ERCC molecules are preferentially sequenced by the Illumina platform (consistent with previous observations^8^) while the ONT MinION platform does not show a preference in this respect. The low GC content ERCC molecules are sequenced equally well in both platforms.

We additionally examined whether the aforementioned deviation is linearly correlated (positively or negatively) with the ERCC transcript length. The comparison between the ONT MinION cDNA counts and Ambion RNA counts showed no statistically significant trend (Fig. 1D). Comparison of the ERCC cDNA abundance derived from the ONT MinION platform with the abundance derived from the Illumina 5’ end fragments showed a statistically significant negative trend (r^2^=0.1, p=0.04) (Fig. 1F). This indicates that the Illumina platform sequences more frequently fragments from short ERCC transcripts. The same is observed when we compared the Illumina 5’ end molecular counts with the Ambion RNA counts (r^2^=0.22, p=0) (Supplementary Fig. S8D). As the tagmentation fragments of the 5’ end from the short ERCC transcripts can be smaller than the ones from the long ERCC transcripts, they are more efficiently sequenced on the Illumina platform.

Including both the GC content and the length of the ERCC transcripts in a component regression model explained 53% of the variation in the deviation (log fold difference) between the observed ERCC cDNA abundance from the ONT MinION platform and the expected ERCC cDNA abundance from the Illumina 5’ end fragments (r^2^=0.53, BIC_score GC+length_ =74 versus BIC_score GC_=82, BIC_score length_ =96).

### The detection of isoforms in a complex cDNA population from the ONT MinION platform complements the Illumina HiSeq 2500 and PacBio RS II platforms

To assess the qualitative and quantitative representation of a complex cDNA population we used cDNA produced from a HEK-293 cell line. We used material from 200 HEK-293 cells in order to reduce the number of different isoforms that can be detected for the same gene, which permits us to detect more genes^14^. We sequenced the same cDNA on an Illumina HiSeq 2500, PacBio RS II and ONT MinION platforms. Initially, we compared the agreement between the long-read sequencing technology platforms PacBio RS II and ONT MinION. Secondly, we compared the agreement of each one of them with the short-read technology Illumina HiSeq 2500. The comparison assessed both the identity and the abundance of the detected isoforms or genes from the ones available in the “NCBI Homo sapiens Annotation Release version 104” transcript database. The ONT MinION run produced 16540 sequenced reads (Table 1). As the amount of cDNA synthesized from the 200 HEK-293 cells was not a lot, the low number of sequenced reads was due to the limited amount of cDNA available to load on the ONT MinION flow cell.

To assess whether in the case of the HEK-293 cDNA library run on the ONT MinION platform, a flow cell specific behavior affected the full length sequencing of the template strand of the cDNA molecules, we introduced into the HEK-293 total RNA, three selected RNA transcripts (“Spike-1”, “Spike-4”, “Spike-7”) at different concentrations. The full length sequencing of the cDNA molecules, from the ONT MinION platform, for these RNA Spikes was then examined. Due to the limited sequencing depth we were able to detect only “Spike-1”. We saw that the template strand of the ONT MinION reads assigned to “Spike-1” was processed overall as full length (Supplementary Fig. S9A, B) and the sequenced length did not change throughout the flow cell run time (Supplementary Fig. S9C).

Initially, we compared the length size distribution of the ONT MinION template reads with the length size distribution of the raw cDNA electrophoresis signal using the Caliper Labchip GX instrument. In Supplementary Fig. S10A we see that both the length size distribution of the ONT MinION template reads and the raw cDNA electrophoresis signal are in good agreement, indicating that the ONT

MinION processed all the cDNA molecules of the complex mRNA population without any specific preference for either the short or the long ones.

After alignment we found 634 isoforms with at least 2 ONT MinION reads assigned to them. In the case of the PacBio RS II platform we detected 6871 isoforms with at least 2 sequenced reads whereas in the case of the Illumina HiSeq 2500 platform we detected 12969 isoforms with an FPKM value of more than 1.

The PacBio ZMW loading procedure, that was used to sequence the HEK-293 cDNA, enriches for molecules longer than 700 bp. Indeed when we compared the overall length size distribution of the PacBio reads, either the longest subread or the high quality Circular Consensus Sequencing reads (CCS reads), with the length size distribution of the raw cDNA electrophoresis signal we saw underrepresentation of molecules lower than 700 bp (Supplementary Fig. S10A).

In Supplementary Fig. S11A we see that the long HEK-293 expressed isoforms detected with the ONT MinION platform (>700 bp in length) are more frequently found as highly expressed among the 10641 isoforms detected with the PacBio RS II platform. Similar picture we take when we compare all the HEK-293 expressed isoforms detected with the ONT MinION platform against the top expressed 10641 isoforms from the Illumina HiSeq 2500. Additionally, the top expressed 400 isoforms of the ONT MinION platform are more frequently found among the highly expressed Illumina HiSeq 2500 isoforms rather than among the highly expressed PacBio RS II isoforms (Supplementary Fig. S11B, D). As the Illumina HiSeq 2500 contains a mixture of short and long isoforms and the short isoforms have usually higher expression than the long ones, more common short isoforms are detected (The FPKM value distribution for isoforms less than 700 bp in length is higher than the one for isoforms more than 700 bp in length).

The example of the RPL41 gene shows the complementary information that the ONT MinION and PacBio RS II platforms can provide (Fig. 2). The ONT MinION provides information for the diversity of both short and long isoforms whereas the PacBio loading procedure followed in this manuscript, can provide information only for the diversity of long isoforms. For example as the Illumina sequenced fragments indicate that for the RPL41 gene the short isoforms are more abundant than the long ones, using the information provided from both platforms is necessary to fully characterize the isoform diversity.

**Figure 2.**
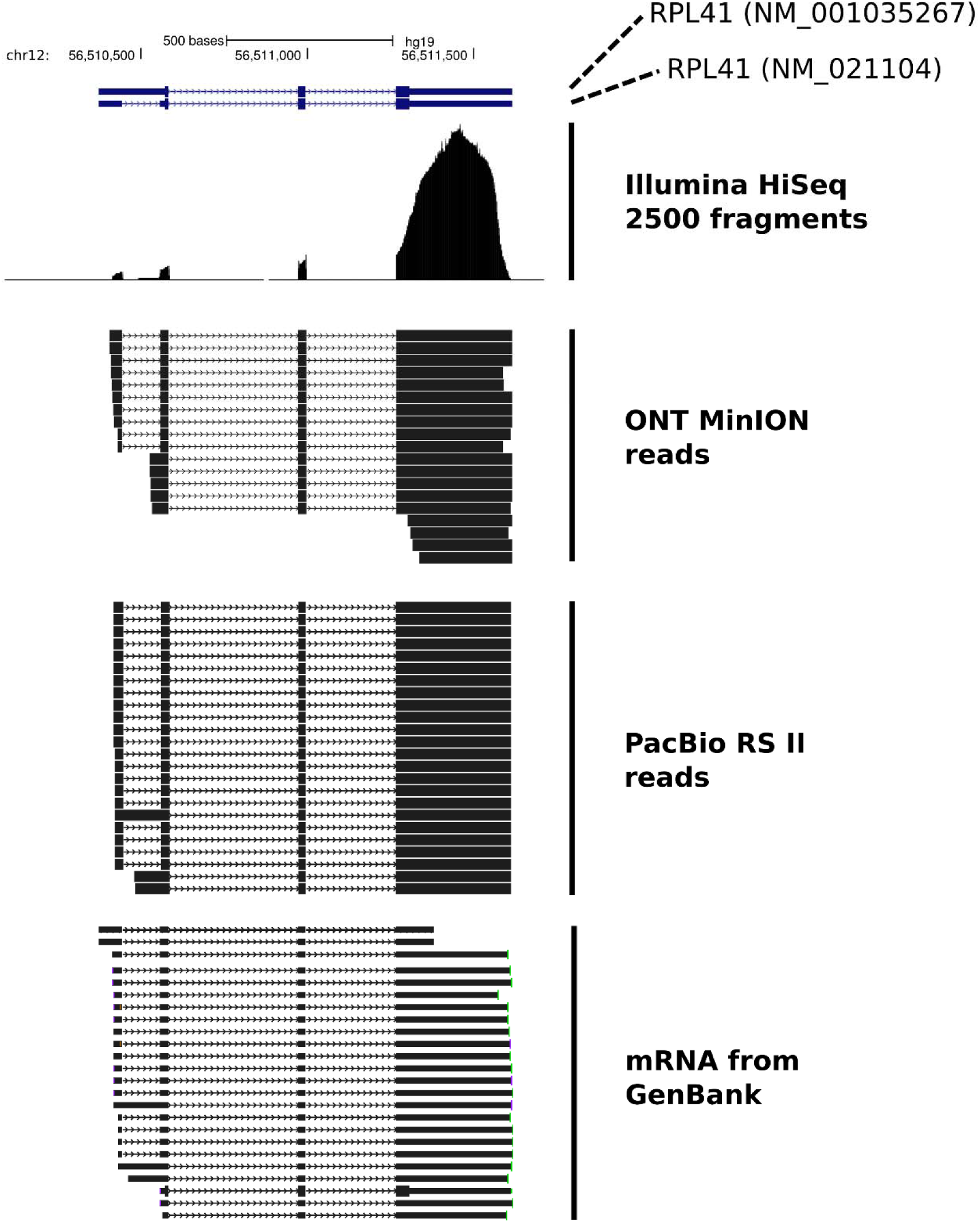
Detection of different isoforms for the RPL41 gene from either the Illumina HiSeq 2500 platform, the ONT MinION platform or the PacBio RS II platform. ONT MinION reads that were sequenced as full length are presented. For the PacBio RS || example the corresponding “Circular Consensus Sequencing” reads are presented. mRNA molecules from GenBank for the RPL41 gene are also shown.

### The expression level estimation of transcripts in a complex mRNA population from the ONT MinION platform is comparable to the Illumina HiSeq 2500 and PacBio RS II platforms

We then assessed the expression level concordance of the isoforms detected from the ONT MinION platform with the expression levels estimated from the Illumina HiSeq 2500 and PacBio RS II platforms. We used the PacBio RS II reads to estimate the abundance of isoforms more than 700 bp in length and the Illumina HiSeq 2500 reads to estimate the abundance of isoforms irrespective of their length. Initially we compared the concordance of the cDNA abundance for the isoforms with length more than 700 bp between the PacBio RS II and Illumina HiSeq 2500 platforms. In Fig. 3A we see a good agreement (r_p_=0.82, r_s_=0.77) between the two platforms. We then compared the ONT MinION with the PacBio RS II and Illumina HiSeq 2500 platforms. In Fig. 3B, C we see that the estimation of the isoform expression levels from the ONT MinION is equally good as the Illumina HiSeq 2500 (r_p_=0.66, r_s_=0.61) and PacBio RS II platforms (r_p_=0.82, r_s_=0.57). We obtained similar results when we compared the expression level concordance at the gene level (Supplementary Fig. S12).

**Figure 3.**
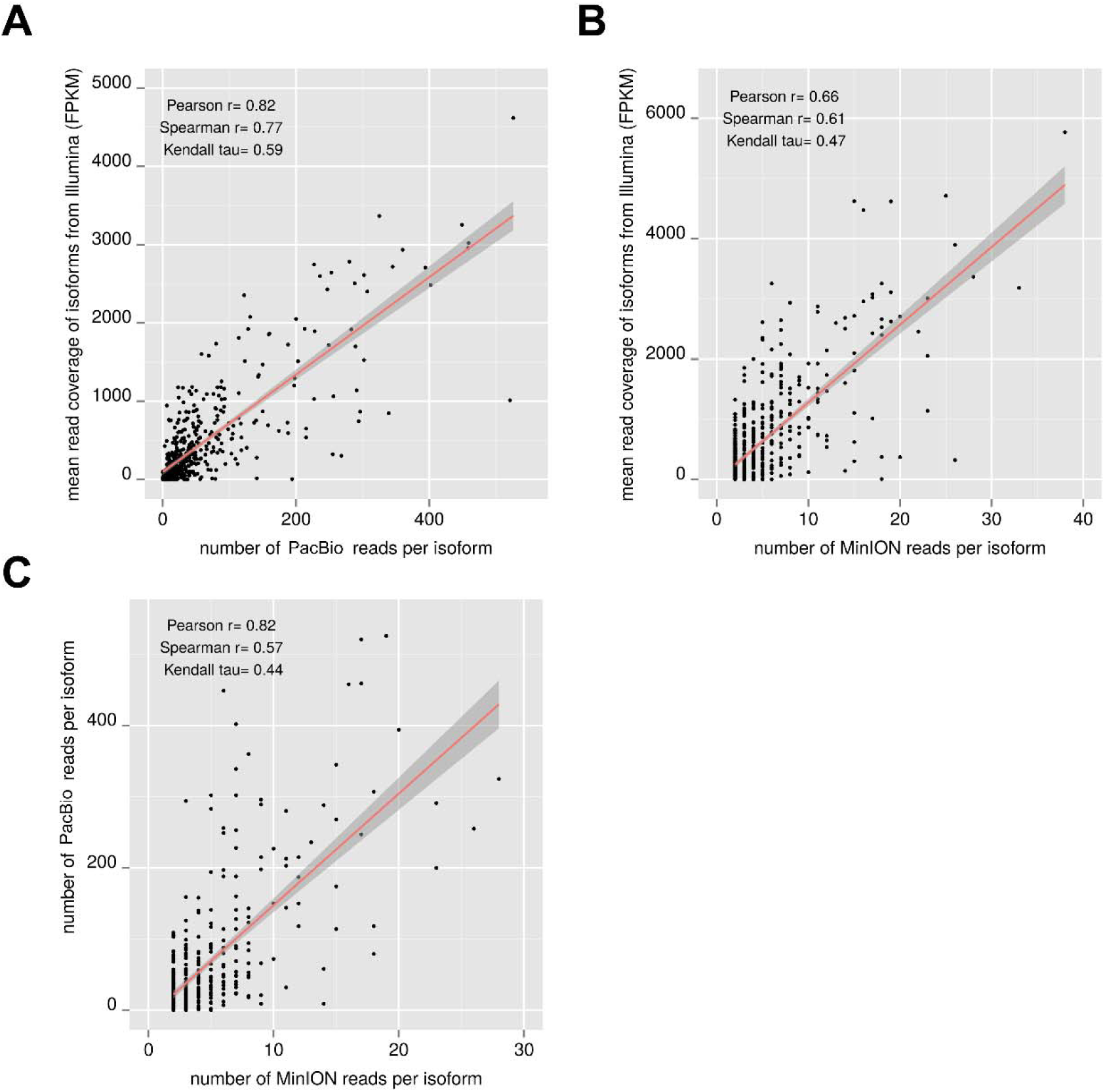
Estimation of the HEK-293 cDNA isoform abundance with three sequencing platforms. The expression level of the HEK-293 isoforms estimated with the ONT MinION platform is compared with the one calculated from either the Illumina HiSeq 2500 (B) or the PacBio RS II platforms (C). For the Illumina HiSeq 2500 platform the expression level is calculated with the FPKM method. For the PacBio RS II or the ONT MinION platforms the counts of sequenced molecules per isoform are presented. The comparison between the Illumina HiSeq 2500 platform and PacBio RS II is also presented (A).

### The TSS and TES of the genes identified with the ONT MinION platform closely agree with the ones identified with the PacBio RS II platform

We compared the position of the transcriptions start site (TSS) and the position of the transcription end site (TES) for the genes detected in both the ONT MinION and PacBio RS II platforms. We selected only genes where we were able to assign at least 3 ONT MinION reads in the case of the TSS comparison or at least 2 MinION reads in the case of the TES comparison. Because the type of isoforms for the same gene detected from the two platforms can be different (Fig. 2), in each gene we assigned as representative of the TSS position agreement the pair of the PacBio RS II and ONT MinION reads with the closest TSS distance between them. Similar approach we used for the TES comparison. In Fig. 4A we see that the TSS of the ONT MinION reads closely follows the TSS of the PacBio RS II reads. Only in seven out of seventy-four genes the distance difference between the TSS is more than 5 base pairs. Similarly, in Fig. 4B we see again that the TES of the ONT MinION reads follows the TES of the PacBio RS II reads. In this case there is a greater variation in the distance difference between the PacBio RS II and ONT MinION reads. In contrast to the PacBio RS II signal the nanopore signal for homopolymer regions (poly-A/poly-T) cannot be adequately resolved in its constituent number of bases. This is why the distance difference of the TES for 26 out of 29 genes is less than or equal to the 30 bp length of the poly-A tail of the cDNA molecules.

**Figure 4.**
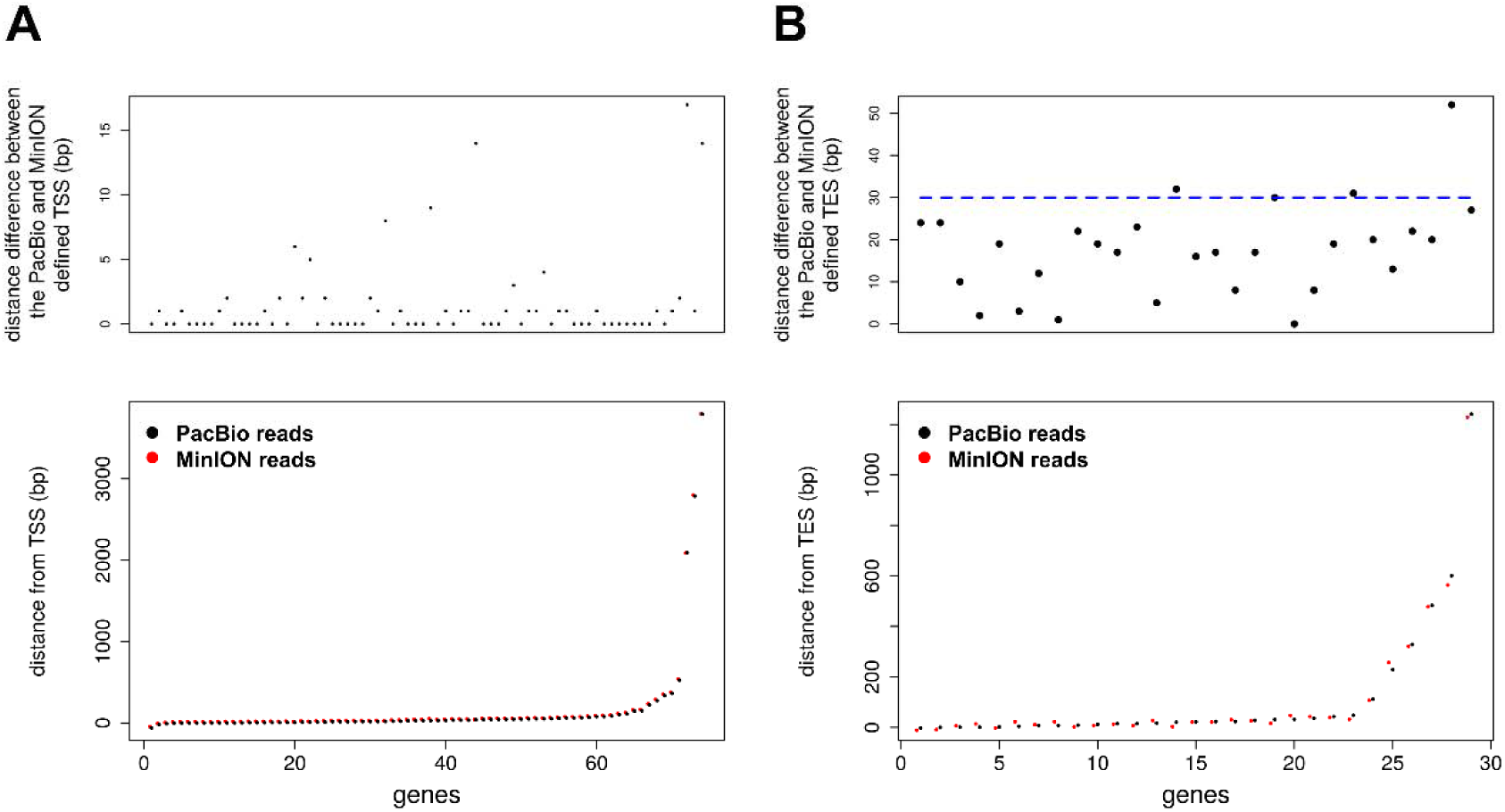
Agreement between the position of the TSS (A) and TES (B) for the genes detected by both the PacBio RS || and the ONT MinION platform. For each gene we assigned as representative of the TSS position agreement, the pair of the PacBio RS II and ONT MinION reads with the closest TSS distance between them. The nucleotide distance difference between the position of the TSS in the two platforms is presented on the top row. In the bottom row the distance from the annotated TSS of the gene isoform, where the pair was assigned to, is presented for each one of the two platforms individually. Similarly for the TES position agreement. The position of the points on the top row correspond to the genes that are present on the same position on the bottom row.

## Discussion

The ONT MinION platform is a third generation/single-molecule sequencing platform that can be used to sequence full length cDNA molecules. Although the reads currently have a higher error rate than other widely used long-read sequencing platforms, in our studies we see that the cDNA molecules are sequenced as full length in the template strand of the sequenced reads. Additionally, the ONT MinION platform does not show any preference in sequencing short or long cDNA molecules from either the simple cDNA population of the ERCC mix or the complex cDNA population of the HEK-293 cells. No bias is also observed for high or low GC content transcripts.

We show that if we use the abundant low quality sequenced reads from the ONT MinION platform, as opposed to only the high quality sequenced reads, we get a consistent result comparable with the other long-read and short-read sequencing platforms. A good agreement between the abundance of targeted cDNA amplicons sequenced with both the ONT MinION and Illumina MiSeq has similarly been observed in another study^10^. Given that the platform can be used for accurate quantification of isoform expression abundance it is very important to be able to use as many sequenced reads as possible with the caveat of being of a lower quality. A combination of the low quality template reads with the high quality 2D reads seems the ideal combination for quantitative accuracy, robustness, reproducibility and maximum usage of the MinION sequenced reads.

As the single-molecule sequencing platforms (PacBio RS II, ONT MinION) do not rely on PCR amplification for sequencing, they usually show lower biases than the short read sequencing platforms^4^. Indeed, in our data we see that the Illumina HiSeq 2500 platform shows an overrepresentation of tagmentation fragments with high GC content or with short length. This is probably due to the PCR amplification biases^6,15^ either at the PCR step after the tagmentation or at the cluster amplification step on the Illumina flow cell^16^.

By introducing commercially synthesized RNA spikes of a known sequence we were able to monitor the performance of the sequencing run, which deems necessary as the ONT MinION platform is still under development. In our case we were able to identify factors that potentially affected the full length sequencing of the cDNA molecules. For example we saw that the increase in the “bias voltage” considerably affected the performance of the full length cDNA sequencing.

The ONT MinION permits the simultaneous quantification of short and long transcripts. On the contrary, the design of the PacBio sequencing flow cell (SMRT Cells) has the disadvantage of biasing towards sequencing short cDNA molecules with the “diffusion loading method” or towards sequencing cDNA molecules longer than 700 bp with the “MagBead loading method”. This can considerably affect the absolute quantification of genes and isoforms as the abundance of short transcripts relative to the long ones cannot be accurately determined.

The ONT MinION platform has been shown to successfully resolve the different isoforms from genes with a complex alternative splicing pattern^10^. Further improvements in the MinION data quality and yield are expected and this will permit the generation of a sufficient number of reads necessary to identify low abundant transcripts but currently the technology is suitable to identify and quantify high abundant cDNA isoforms. cDNA normalization methods based on duplex-specific nuclease (DSN)^17^, aimed at enhancing the detection of rare transcripts in eukaryotic cDNA libraries by decreasing the prevalence of highly abundant transcripts, can help increase the resolution in the median to low abundant transcript range at the current ONT MinION sequencing yield.

## Methods

### Materials

The following materials were used: “ERCC RNA Spike-In Mix 1” stock (#4456740, Ambion, Thermo Fisher), RNA storage solution (Ambion, Thermo Fisher), Triton-X 100 (molecular biology grade, Sigma), dNTP mix (Advantage UltraPure PCR Deoxynucleotide Mix 10 mM each dNTP, Clontech), RNAse inhibitor (Clontech), Superscript III enzyme and Superscript III first-strand buffer (Invitrogen, Thermo Fisher), ultrapure DTT (Invitrogen, Thermo Fisher), Betaine (Sigma), MgCl_2_ (Sigma), oligo-dT30V primer (IDT), template switch primer (IDT), cDNA amplification primer (IDT) (the sequences are given in Supplementary Fig. S15), 2X KAPA2G Robust HotStart ReadyMix (kapabiosystems), AMPure XP beads (Beckman Coulter), NEBNext End Repair Module (New England Biolabs, UK), NEBNext dA-Tailing Module (New England Biolabs, UK).

### ERCC cDNA production

An adaptation of the Smart-seq2 method ^18^ was used to produce full length cDNA molecules from the ERCC RNA transcripts. Two different ERCC RNA concentrations were used mimicking conditions where the poly-A RNA was derived from either a bulk of cells or from a few cells (~200 cells / single cell). In the first case, the “ERCC RNA Spike-In Mix 1” stock (32 ngs/ul) was diluted 10 times in RNA storage solution down to 3.2 ngs/ul. In the second case, the “ERCC RNA Spike-In Mix 1” stock was diluted 500 times in RNA storage solution down to 0.064 ngs/ul. The poly-A hybridization step, of the first strand cDNA synthesis primer, was performed in a 3 ul volume reaction (3.2 ngs or 0.064 ngs ERCC RNA Spike-In, 0.12% Triton-X 100, 4.2 μM of oligo-dT30V, 2.8 mM dNTP mix, 1X Superscript III first-strand buffer, 20 U RNAse inhibitor) with the following thermocycler parameters 72°C for 3 min, 4°C for 10 min, 25°C for 1 min. The synthesis of the first and second strand of cDNA was performed by topping the 3 ul reaction up to a 7 ul volume reaction (final concentration: 1X Superscript III first-strand buffer, 2.5 mM DTT, 1.2 μM template switch primer, 27 U RNAse inhibitor, 70 U SuperScript III reverse transcriptase, 0.5 M betaine, 3.6 mM MgCl_2_). The cDNA synthesis was performed with the following thermocycling parameters: 1 cycle of [50 °C for 90 min], 10 cycles of [55 °C for 2 min, 50 °C for 2 min], 5 cycles of [60 °C for 2 min, 55 °C for 2 min], 1 cycle of [70 °C for 15 min]. Afterwards the double stranded cDNA in the 7 ul reaction was amplified in a 70 ul PCR reaction containing 12 μM cDNA amplification primers and 1x KAPA2G Robust HotStart ReadyMix. Depending on the initial amount of RNA in the reaction we used 14 cycles of amplification for 3.2 ngs of RNA or 21 cycles of amplification for 0.064 ngs of RNA. In the case of 14 cycles of cDNA amplification the cDNA amplification protocol used was the following: 1 cycle of [95°C for 1 min], 5 cycles of [95°C for 20 seconds, 58°C for 4 minutes, 68°C for 6 minutes], 9 cycles of [95°C for 20 seconds, 64°C for 30 seconds, 68°C for 6 minutes], 1 cycle of [72°C for 10 minutes]. In case of 21 cycles of cDNA amplification the cDNA amplification protocol used was the following: 1 cycle of [95°C for 1 min], 5 cycles of [95°C for 20 seconds, 58°C for 4 minutes, 68°C for 6 minutes], 9 cycles of [95°C for 20 seconds, 64°C for 30 seconds, 68°C for 6 minutes], 7 cycles of [95°C for 30 seconds, 64°C for 30 seconds, 68°C for 7 minutes], 1 cycle of [72°C for 10 minutes]. The amplified product was subsequently cleaned with 0.9X sample volume AMPure XP beads and eluted in H_2_0 for a total of 2400 ngs of cDNA per 3.2 ngs of initial RNA material (or 5400 ngs of cDNA per 0.064 ngs of initial RNA material). For sequencing on the Illumina HiSeq 2500 or MiSeq platforms, the cDNA was tagmented using the Nextera XT DNA Library Preparation Kit (Illumina Inc) as described in supplementary methods.

### ONT MinION sequencing of the ERCC cDNA population

The full length cDNA libraries were sequenced with version r7 and version r7.3 MinION flow cell. The ONT MinION Genomic DNA Sequencing Kit reagents (SQK-MAP0003 for the r7 flow cell, SQK-MAP0004 or SQK-MAP0005 for the r7.3 flow cells as indicated on Table 1) and the 2D cDNA sequencing protocol were used to prepare the libraries. The amount of starting material used is presented in Supplementary Table S2. In each case the cDNA was end-repaired in a 100 ul reaction containing: 0.45-2 ug cDNA (depending on the run) in a 80 ul solution, 10 ul of 10X End-repair buffer (from the NEBNext End Repair Module), 5ul End-repair enzyme mix (from the NEBNext End Repair Module) and 5 ul of nuclease-free water. In the case where we started with 3 ug of cDNA, two reactions with 1.5 ug of cDNA material were used (ERCC experiment 4, Supplementary Table S2). The end-repair reaction was then incubated at 25°C in a thermocycler for 30 minutes. Afterwards we added 1X sample volume Ampure XP beads to the End-Repair reaction. The solution was mixed by pipetting and the DNA was allowed to bind to the beads by rotating for 5 minutes on a HulaMixer (Thermo Fisher). Then the beads were pelleted on a magnet, the supernatant was aspired off and the beads, while they stayed on the magnet, were washed twice with 200 μl of freshly prepared 70% ethanol. The tube was centrifuged to collect residual liquid at the bottom and the residual wash solution was aspirated. The DNA was eluted from the beads by resuspending the beads in 26 ul of 10 mM Tris-HCl pH 8.5 and the beads were incubated for 5 minutes at room temperature. The isolated DNA was quantified on the Qubit fluorimeter (Thermo Fisher). For the subsequent dA tailing step all the eluted DNA was used in the following reaction: cDNA library solution from the previous step (>750 ng), 3 ul 10x dA-tailing buffer (from the NEBNext dA-Tailing Module), 2 ul dA-tailing enzyme (from the NEBNext dA-Tailing Module). Subsequently, depending on the protocol version the DNA was cleaned with the Ampure XP beads as described above (SQK-MAP0005 kit version) or used immediately in the ligation reaction (SQK-MAP0004, SQK-MAP0003 kit version). In the ligation reaction the DNA was mixed with the following reagents: 30 ul dA-tailed DNA, 10 ul Adapter Mix, 10 ul HP adapter, 50 ul Blunt/TA ligase Master Mix. The reaction was incubated at room temperature for 10 minutes. Depending on the protocol the cDNA library was either loaded on the flow cell as is (SQK-MAP0003 kit version) or enriched for sequences that bear the hairpin (SQK-MAP0004, SQK-MAP0005 kit version) as follows.10 ul of His-beads (Dynabeads His-Tag, Thermo Fisher) were washed two times with 1x Bead Binding Buffer (SQK-MAP0005 kit), then resuspended in 100 μl of the 2x Bead Binding Buffer (SQK-MAP0005 kit) and added to the adaptor ligated DNA. The bead bound DNA was then washed two times with 200 μl 1x Bead Binding Buffer (SQK-MAP0005 kit) and eluted with Elution Buffer (SQK-MAP0005 kit) with the amount presented in Supplementary Table S2. The parameters used to align the ONT MinION reads are presented in supplementary methods. Considerations on the amount of cDNA library to adaptor molecules used are also presented in supplementary methods. The number of aligned reads per ERCC transcript for the different MinION experiments is provided in the supplementary results excel file.

### HEK-293 cDNA production and ONT MinION sequencing

cDNA was produced from 200 HEK-293 cells expressing shRNAs targeting PBRM1 (the cells were kindly provided from Dr. Yasser Riazalhosseini). The HEK-293 cDNA production was based on the Single-Cell cDNA Library preparation protocol for mRNA Sequencing for the Fluidigm C1 machine as described in detail in supplementary methods. The HEK-293 cDNA library was sequenced with a version r7.3 MinION flow cell. The ONT MinION Genomic DNA Sequencing Kit reagents (SQK-MAP0005) and the 2D cDNA sequencing protocol were used to prepare the libraries. The library preparation method was performed similarly to the ERCC cDNA and is presented in the supplementary methods.

### PacBio RS II library preparation and sequencing of the HEK-293 cDNA

The amplification product from the cDNA preparation was prepared for sequencing using 28ng of input material. The protocol we used^19^ is a modified version of the PacBio RS II library preparation (SMRTbell template prep Kit). First a DNA damage repair step and end repair step is performed followed by purification (AMPure XP beads). The hairpin is then ligated to the fragments. Here 500ng of carrier DNA (Clontech pBR322) is added to the library before exonuclease treatment and additional purification. Following this a sequencing primer with a poly-A tail is hybridized to the hairpin ligated to the fragments. Magnetic dT beads are then used to both separate the library from the carrier and also to load the fragments onto the SMRT cells. The parameters used to align the PacBio RS II reads are presented in supplementary methods.

### Data access

All sequencing datasets have been submitted to the ENA database under the accession number PRJEB11818. A description of each file in the ENA archive is provided in the supplementary excel file.

## Acknowledgements

We thank Dr. Timothée Revil and Louis Letourneau for useful discussion during the experiments and the analysis. We also thank Pierre Bérubé, Geneviève Geneau, Patrick Willet and Dr. Alfredo Staffa for the PacBio library preparation and sequencing. The work was supported by Genome Canada Science Technology Innovation Centre grant, the Compute Canada Resource Allocation Project wst-164-ab and the Genome Innovation Node grants to Dr. Jiannis Ragoussis.

## Author Contributions

Performed experiments: S.O., Y.C.W. Analyzed data: S.O., Y.C.W., H.D., D.B. Generated Fig.s and tables: S.O. Wrote manuscript: S.O., J.R. All authors critically reviewed the manuscript.

## Competing interests

J.R. is a member of the Oxford Nanopore Technologies Inc. MinION Access Programme and the MinION instrument and R7, R7.3 flowcells were received free of charge.

